# Structured navigation in a goal-directed task reveals flexible spatial coding

**DOI:** 10.64898/2026.07.27.741044

**Authors:** Tucker G Fisher, Marielena Sosa, Alexander Gonzalez, Xiaotong Cheng, Lisa M Giocomo

## Abstract

Animal survival depends on accurate navigation. The last several decades have revealed that neurons encoding position and orientation can support navigation through the construction of an internal map of space for the external world. However, current experimental approaches for studying this system require tradeoffs between behavioral richness and experimental control; open field tasks often lack structured trials or trajectories and virtual reality paradigms constrain natural movement variables. Here, we introduce a platform that combines freely moving behavior with control over environmental stimuli, trajectories, and reward locations. This platform is compatible with chronic Neuropixels recordings, dynamic optical projection of visual stimuli and real-time pose estimation. Using this platform, we show rats can learn a multistage spatial targeting task and that high density Neuropixels recordings can be performed in conjunction. Recordings in medial entorhinal cortex revealed canonical spatial and head direction tuning, along with population-level activity that tracked distance to task-relevant goals. Together, this platform provides a powerful approach for examining the neural basis of flexible navigation under experimentally controlled yet naturalistic conditions.

## Introduction

Many species, including insects, fish, rodents, bats, non-human primates, and humans, rely on neurons that encode position, orientation, speed, and landmarks to construct an internal representation of external space (Yartsev et al. 2011; Killian et al. 2012; Jacobs et al. 2013; Kim et al. 2017; Yang et al. 2024; Hafting et al. 2005). In mammals, this spatial system has been extensively studied in the medial entorhinal cortex (MEC), a component of the entorhinal-hippocampal circuit (Rowland et al. 2016). MEC neurons are defined by their firing patterns observed as animals navigate and include cells for encoding position (grid cells and spatial cells with non-periodic firing), landmarks (border and object vector cells) and velocity (head direction and speed cells) (Sargolini et al. 2006; Hafting et al. 2005; Solstad et al. 2008; Høydal et al. 2019; Kropff et al. 2015; Diehl et al. 2017). Together, these neurons are hypothesized to provide a neural substrate for generating an internal spatial map capable of supporting navigation and memory (Buzsáki and Moser 2013; Hartley et al. 2014; McNaughton et al. 2006).

Much of the work on spatial coding in the MEC has relied on a handful of paradigms, many of which involve rodents foraging for randomly scattered food rewards in an open field (Hafting et al. 2005; Solstad et al. 2008; Kropff et al. 2015; Diehl et al. 2017). Ethologically relevant behavior however, tends to be highly dynamic and complex, requiring animals to flexibly adapt their trajectories based on changes to their goals, the location of rewards and their sensory environments (McMillan and Kaufman 1995; Benhamou 1991; Thompson 1982). Moreover, while open field tasks allow naturalistic locomotion and broad sampling of spatial variables, they lack repeated trajectories, defined decision points and trial structure. As a result, aligning neural activity to changing behavioral epochs or goals remains challenging, and spatial locations may only be sampled once in each behavioral session, limiting statistical power. At the other extreme, head-fixed virtual reality paradigms provide precise control over stimuli and enable repeated trajectories (Dombeck et al. 2010; Harvey et al. 2009). However, this approach constrains natural movement, dampens vestibular feedback and restricts the range of behaviors the animal can naturally express (Pak et al. 2026). Thus, current approaches impose a tradeoff between behavioral richness and experimental control.

Addressing this challenge is timely given recent evidence that MEC spatial representations are state-dependent. MEC grid, border and non-grid spatial cells undergo substantial changes in their firing field shapes, locations and firing rates when navigation incorporates remembered reward locations compared to random foraging (Butler et al. 2019; Boccara et al. 2019). In parallel, rapid technological developments now allow unprecedented access to recordings of large populations of MEC neurons (Steinmetz et al. 2021). These recent developments point to a key need for a broader repertoire of behavioral paradigms and platforms that are compatible with state-of-the-art electrophysiological techniques for studying navigation.

Here, we introduce a behavioral and hardware platform to bridge this gap. Our goal was to combine the richness of freely moving behavior in an open field with the precise control over stimuli, trajectories, and reward contingencies characteristic of virtual reality setups. To do this, we designed a large, stable arena compatible with chronic Neuropixels recordings, optical projection of dynamic visual targets, real-time pose estimation, and modular task control. To validate this system, we designed a spatial targeting task that supports repeated, structured trajectories without physically constraining the movement of the animal. Using this setup, we show that MEC neural recordings capture both canonical position and head direction tuning, as well as population-level signals related to distance to task-relevant goal locations. Thus, this platform provides a framework for examining how neural maps support flexible, goal-directed navigation in dynamic environments.

## Results

### Freely moving behavior rig design

Here, we endeavored to combine the precise control over stimuli and repeatable trajectories characteristic of head-fixed virtual reality paradigms with the richness of freely moving behavior in a physical open field environment. While traditional open field tasks allow animals to express a broad range of locomotor gaits and head directions, trajectories are rarely repeated. This lack of repetition presents challenges for statistical rigor, as certain positions may only be sampled once. Additionally, the commonly used random foraging paradigm makes it difficult to align specific behavioral states or goals with defined epochs of neural activity. Our goal was therefore to develop a system for rodents that preserves the key features of unrestricted movement while providing more experimental control over task structure, trajectory repetition, positions sampled, and reward contingencies.

Central to the design is a large, mechanically stable arena, which was optimized for optical projection of targets onto the arena floor and compatibility with chronic electrophysiological neural recordings (Figure 1A-C). The arena frame was constructed from 80/20 aluminum. The floor consisted of a single sheet of acrylic with a satin finish; this provided a durable floor that also facilitated optical bottom-up projection of visual stimuli. The arena had four translucent wells for delivering liquid rewards. Each well was set within the rectangular arena so that the lick port was ∼9 cm from the wall (Figure 1D, wells shown in black to highlight them in the image). The arena walls were constructed from pieces of opaque black satin acrylic and were fixed to the frame of the arena by magnets; this facilitated easy removal for cleaning and maintenance of the environment. An overhead camera and wide-angle perspective correcting rectilinear lens, to minimize distortion across the arena, were affixed to a cross bar above the arena. The slip ring and recording cable suspension system were affixed to a second upper cross bar.

**Figure 1.**
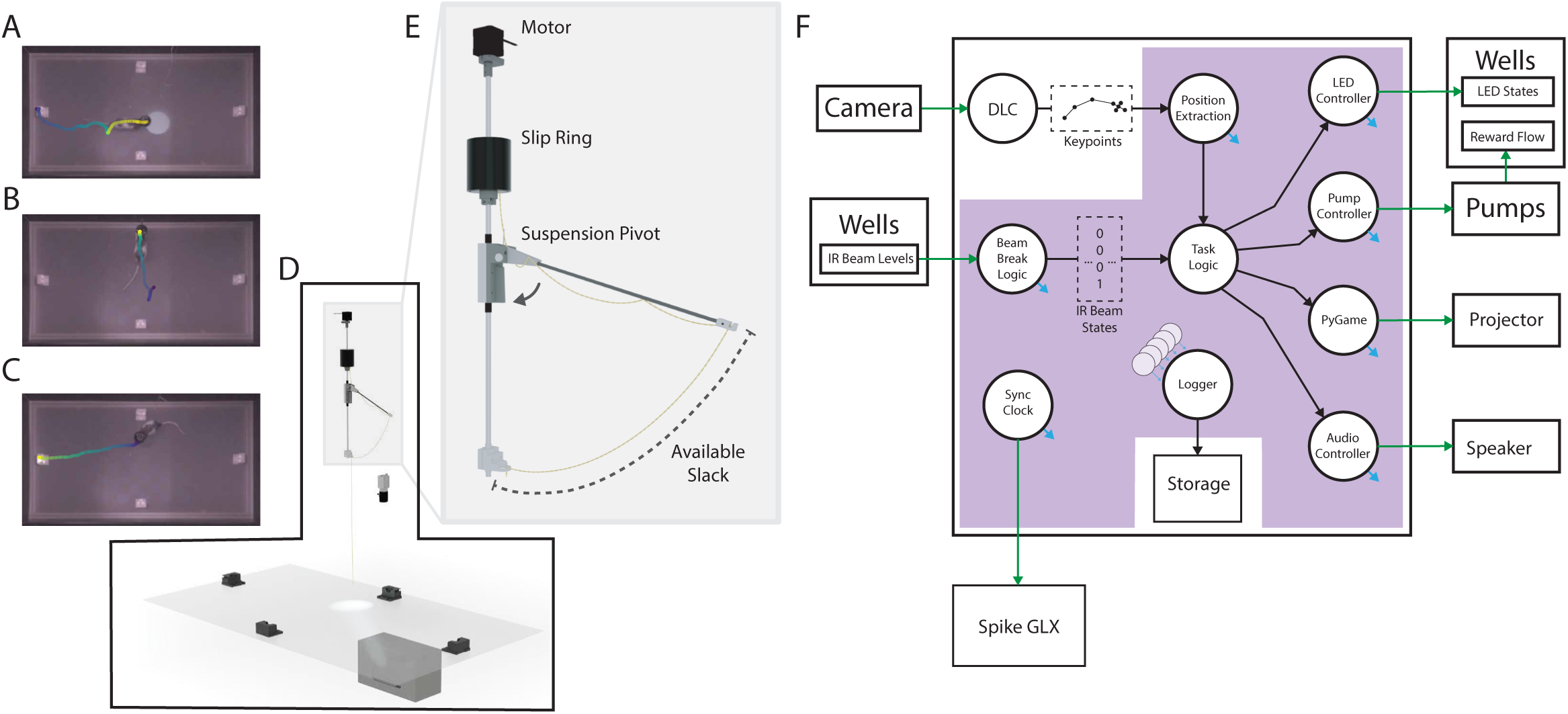
Overview of the behavioral arena, task and hardware for task implementation and data collection. A) Overhead view of the arena. Example past trajectory in which the animal has made a run to collide with the illuminated target. The blue to yellow gradient indicates the animal’s position along this trajectory over time. B) Example past trajectory in which the animal has made a run to collect a reward at a well. Color coded as in (A). C) Example future trajectory in which the animal is about to make a run to initiate the next trial. Wells can be illuminated to cue the animal thar a reward is guaranteed at that location. Color coded as in (A). D) Three-dimensional render of key behavioral rig components. Wells for reward delivery are arrayed offset from the walls of the arena (walls not shown) and a short-throw projector projects visual stimuli onto the translucent acrylic floor. A camera overhead records the session. In addition, frames captured are used to determine the animal’s position in the arena during the task. E) Three-dimensional render of the custom slip ring and suspension system. In this illustration, the suspension arm is in its highest position, with the cable slack fully retracted. When the animal visits the corner of the arena and requires more cable length, the arm is pulled down to a more acute angle. The whole assembly below the slip ring is free to rotate, which prevents coiling of the Neuropixels cable (in yellow) and the fishing line tether (not shown). A motor coupled to the central rod drives rotation. F) Signal flow through the experimental setup, highlighting the core elements needed to run the task. Rectangles represent individual hardware components. The central rectangle represents the task computer. Circles represent processes such as DeepLabCut Live pose extraction. Processes within the purple background comprise the Linux Comodular Realtime Interactive Computation Engine (LiCoRICE) modules. Processes with blue arrows emit signals to the logger module to be stored. Green arrows indicate signal flow between separate hardware components.

Visual input to the environment was generated by a short-throw projector, mounted under the arena, which projected visual cues onto the underside of the acrylic floor (Figure 1D, bottom). This produced a clear and spatially concise visual cue that could move to a different position on each trial. Auditory tones were delivered via a mono-speaker mounted below the arena (not shown). Both the visual and auditory inputs were controlled by the task software, allowing dynamic yet temporally precise cue control.

Chronic electrophysiology in freely moving animals requires rotational freedom while limiting the accumulation of twisting and cable slack. To achieve this, we designed a motorized slip ring that allowed for uncoiling of the Neuropixels acquisition cable without interrupting the recording. With this system, the rats could turn freely while performing the task. An additional challenge is the large size and shape of the environment, such that a static cable length is insufficient to both reach the corner of the environment and avoid the accumulation of excessive slack as the rat moved through the center of the arena. To manage this, we designed a swinging arm suspension system that prevented a distracting cable loop from being within reach of the rat, while retaining stable and smooth action even during slip ring rotation (Figure 1E). Our design leveraged a swing arm and spring chosen to offset the weight of the extra cable and headstage, allowing the animals to reach the edges of the environment with minimal effort.

### Task Related Hardware and Software Infrastructure

All task related signals were managed by a central task computer. From the perspective of this task machine, each hardware element was a “source”, providing information about the state of the rat within the environment or a “sink”, providing an endpoint where the task extends a signal into the environment (Figure 1F, green arrows going into and out of the task computer, represented as a central box). Each element was managed by a single software module allowing for a clear and compositional task design supporting flexible simplification or expansion of the system.

An accurate estimate of the moment-to-moment position of the animal is central to controlling this task. Pose estimation was carried out on frames from the overhead camera by a DeepLabCut Live model. We specifically employed a MobileNet architecture during task time to achieve inference times exceeding the frame capture rate, ensuring the pose estimates did not lag changes in behavior.

Each reward well was equipped with an infrared (IR) emitter and receiver pair. A drop in IR light indicated the beam was broken; the task software treated this as a nose-poke, or “reporting to a well.” In early pre-training phases, animals occasionally broke the beam while stepping on or near the well, but this behavior was almost entirely extinguished with training. An internal light emitting diode (LED) in each well was used during training and in certain phases of the task to provide an explicit cue to the animal that the well was active. At each well, reward was provided through a tube connecting the well inlet to a syringe pump located away from the recording area; this placement minimized electrical interference.

All software required for the task, with the exception of DeepLabCut Live inference, ran as a model under the Linux Comodular Realtime Interactive Computation Engine (LiCoRICE) (Mehrotra et al. 2018). Models are organized as blocks of Python code that each have a specific responsibility over internal signals (Figure 1F, circles within the purple background indicate LiCoRICE modules or logical groupings of related modules). Once properly configured, LiCoRICE manages translating these modules into Cython, compiling them, and coordinating them as a real-time or near-real-time model. In our case, the LiCoRICE model was configured to update its state on a precise 10 ms interval.

### Task structure and overview

To validate the hardware integration and demonstrate the strengths of our experimental apparatus, we designed a multistage spatial targeting task that combined freely-moving navigation with a clear trial structure. Each “block” consisted of multiple trials. Each trial in this task had three distinct phases: targeting, reporting, and pre-initiation. In the first phase, “targeting”, animals were required to locate and intersect their head with an unrewarded target location (i.e. visual cue) projected onto the floor (Figure 2A, top panels of Block A or B). Following a hold period where the rat was required to keep its head within the boundaries of the target location, the “reporting” phase began. The remembered reward well became active, indicated by an affirmative tone and the disappearance of the visual cue denoting the target location. The remembered reward well location was always consistent within a block. Each block began with a hint trial, in which an LED within the appropriate reward well blinked to instruct the animal if the block was “A” (rewarded at well 1) or “B” (rewarded at well 3). On subsequent trials the hint was successively delayed, requiring animals to progress on memory or wait an extended period of time for a cue (Figure 2A, middle panels of Block A or B). Animals performed “memory trajectories” when navigating from target to reward well without hints. The final phase, “pre-initiation”, was an intermediate phase between trials where the rat was instructed to reset its position to one of the two ends of the environment and initiate the next trial (Figure 2A, bottom panels of Block A or B). After reward consumption, at the end of the report phase and 1.5 second delay, one of the wells at the end of the environment became active, indicated by the internal LED. A new trial was self-initiated by the rat once it broke the IR beam and consumed a small initiation reward.

**Figure 2.**
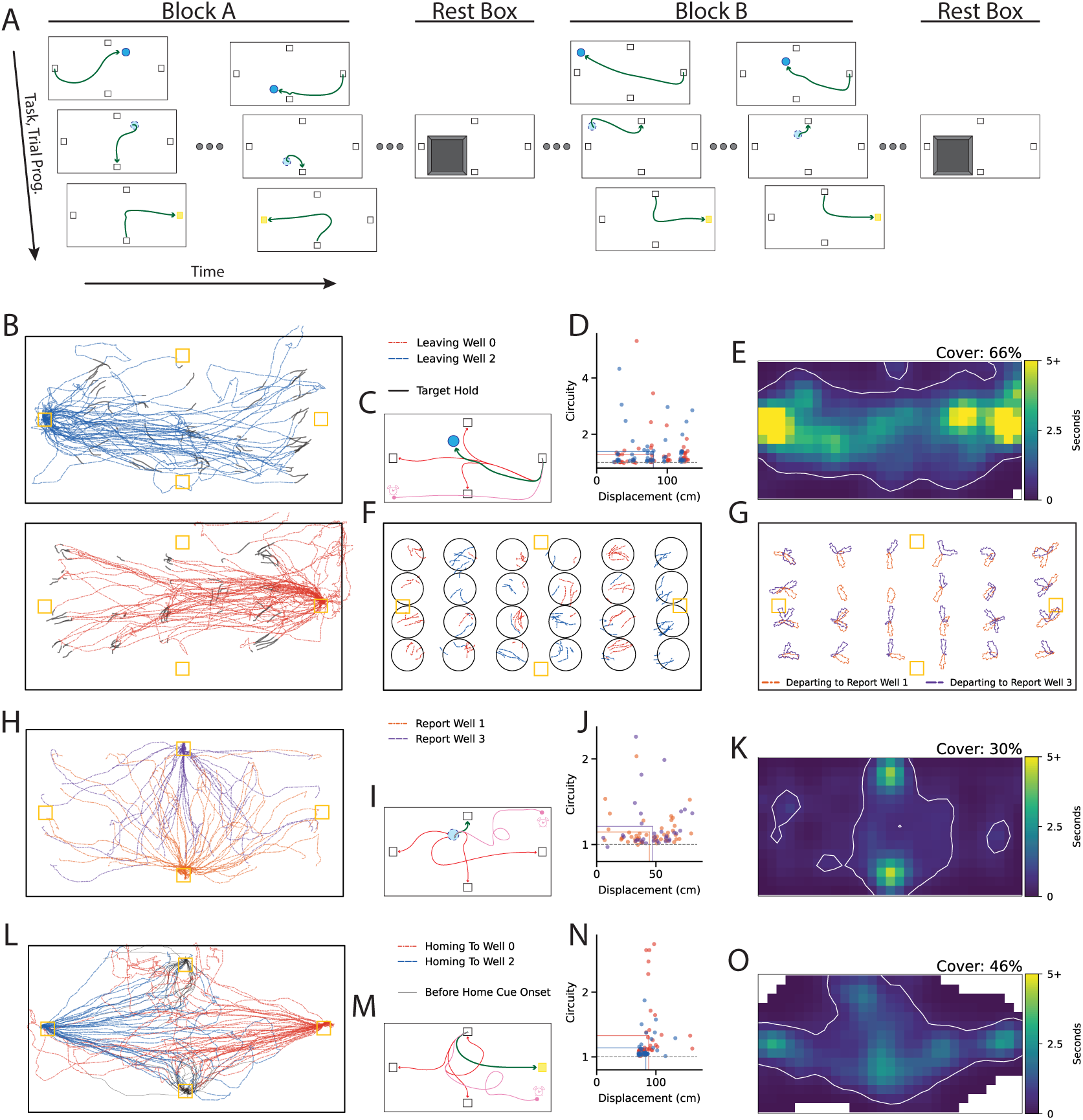
Task structure and behavioral performance on a goal-directed navigation task. A) Schematic of the task progression over time, with offset columns representing the flow of a single trial. Blue circles indicate targets. Green lines indicate valid example trajectories within each phase. Black rectangles indicate well positions, wells on the left (well 2) and right (well 0) are for trial initiation, and wells at the top (well 3) and the bottom (well 1) are choices for the report phase. Light blue dotted circles indicate the disappearance of the target. Target disappearance signaled that the required target hold is complete and the remembered reward location was now active. Yellow squares indicate wells with the internal Light Emitting Diode (LED) on. These LEDs instructed the animal where to initiate a trial. Animals performed trials in blocks. Within a block the reward location, either top or bottom, remained consistent. The arenas with a grey square, the rest box, represent rest periods between blocks of trials. B) All target phase trajectories (blue) initiated at well 2 and trajectories (red) initiated at well 0 on an example day. The surrounding black rectangle represents where the walls meet the floor of the arena. The tracked points outside of the floor boundary rectangle are a result of the animal rearing against the walls of the arena. Yellow squares indicate the four well locations. Thin grey lines extend the target trajectories into the target hold phase, when the animal was colliding with the target. C) Schematic of possible error states for the task phase behavior associated with (B). Green lines indicate the correct trajectory that the animal should have taken to the target (blue circle). The animal may instead make an invalid visit to a reward well (red lines) or fail to transition to the next state after some time period resulting in the task timing out (pink line). D) Circuity, or route directness, for each trajectory in (B). Vertical and horizontal lines show average displacement and circuity, respectively. The grey dotted line is at circuity of one. Colors correspond with (B). E) Smoothed occupancy map in 5x5 cm spatial bins, color coded for high (yellow) versus low (blue) occupancy (Butler et al. 2019). White lines surround areas with greater than 500 ms of data per bin. Coverage (cover) indicates the percent of bins with greater than 500 ms occupancy. White bins have no occupancy. Data are combined from example trajectories in (B). F) Target hold phase colored by initiation well with common elements coded as in (B). Black circles represent possible target locations. Trajectories extracted by the offline DeepLabCut model are slightly different than those extracted by the DeepLabCut model during experimental sessions, resulting in the apparent overflow of target boundaries. G) Head direction occupancy maps for trajectories departing from each target location colored for endpoint well. Maps are smoothed and normalized to the maximum bin. H) Report phase, post-target memory guided trajectories colored by the rewarded well (purple versus orange). Other common elements coded as in (B). I) Schematic of possible errors associated with reporting to the remembered reward location after target collision and hold. Coded as in (C). J) Circuity for each trajectory in (H). Vertical and horizontal lines show average displacement and circuity. K) As in (E) but for trajectories in (H). L) Pre-initiation phase, trajectories post-consumption of reward to the next trial initiation, colored by initiation well. Black lines indicate positions traversed before the initiation well cue. Trajectories and other common elements color coded as in (B). M) Schematic of possible errors associated with initiating a new trial after reward. Coded as in (C) with an additional yellow rectangle indicating a well 2 LED cue. N) Circuity for each trajectory in (L). Vertical and horizontal lines show average displacement and circuity. O) As in (E,K) but for trajectories in (L).

Two types of error states encouraged animals to remain on task. First, if an animal visited an inactive well and broke the IR beam, this triggered a “beam break error”. Second, if an animal did not complete a phase within the time allowed, this triggered a “timing out error”.

The task was designed to encourage repeated, structured navigational behavior without physical constraints. Spatial coverage is a common task requirement for the purpose of characterizing spatial cells, a process that at a minimum requires the animal to be in many different locations in the arena and in each location for a sufficient duration of time (O’Keefe and Dostrovsky 1971; Langston et al. 2010). For this reason, we exploited the ability to force exploration, instructing the animals to collide with targets chosen from a regular grid spanning the environment.

In the targeting phase, the animal had to collide with the light pool, or target, directly, from either initiation well—resulting in bundles of trajectories that share a common origin and endpoint (Figure 2B, S1C). In this phase all wells are considered inactive, meaning that breaking any well’s IR beam resulted in a beam break error and if the phase was not completed in under 30 seconds, the task would time out (Figure 2C). Trajectories with the same origin and endpoint clustered together and, besides a few outliers, the majority of trajectories had low circuity indicating they were direct to target (Figure 2D). Within a block, target locations were chosen at random, excluding the column of target locations next to the initiation well, without replacement until all target locations were exhausted at which point target locations were redrawn. This promoted an even distribution of targeting trajectories and target holds, resulting in good coverage of the environment (Figure 2E,F).

In the reporting phase, animals were first tasked with making uncued, memory guided trajectories to a remembered reward well, after learning the reward well through a hint LED at the start of a block. Head direction summaries of the memory trajectories from each target to reward combination highlight the ballistic nature of the behavior during this phase, noting that a brief pause before departing the target zone was common (Figure 2G,H, only memory trials shown). During the reporting phase, all wells except for the remembered reward well are inactive, and the task will time out if the rat does not report within 30 seconds (Figure 2I). Again, the trajectories had a circuity clustered near one (Figure 2J). During this phase good spatial occupancy occurs where the trajectories come together in the middle of the arena (Figure 2K).

In the pre-initiation phase, the animal is cued by LED to collect a smaller initiation reward at one of the initiation wells (Figure 2L). Once the initiation well is active, visiting any other well triggers a beam break error and if the phase was not completed in under 20 seconds the task would time out (Figure 2M). The trajectories in this phase had less displacement variability than the other two phases (Figure 2N). Coverage in this phase was also highest in the center of the arena (Figure 2O).

### Behavioral performance across sessions

The behavioral trends in the example session are consistent with the trends across all animals and days. Coverage was high in the target phase. The report and pre-initiation phases had low coverage. Further analysis of report and pre-initiation trajectories showed better coverage in the middle of the arena: excluding animal J, each phase had an average coverage of around 48%, between 10-96% for pre-init and 10-86% for report, within a centered square with 45cm side lengths (Supplemental Figure 1). Overall except for animal J, the weakest performer, error-free memory trial coverage was between 72% and 88% with an average of 80% coverage for all recording days (Figure 3A). These coverage percentages are consistent with what has been routinely used as a criteria for classifying spatial MEC cells in prior works (Langston et al. 2010; Wills et al. 2010). It is important to note that achieving these coverage percentages required combining blocks within a day.

**Figure 3:**
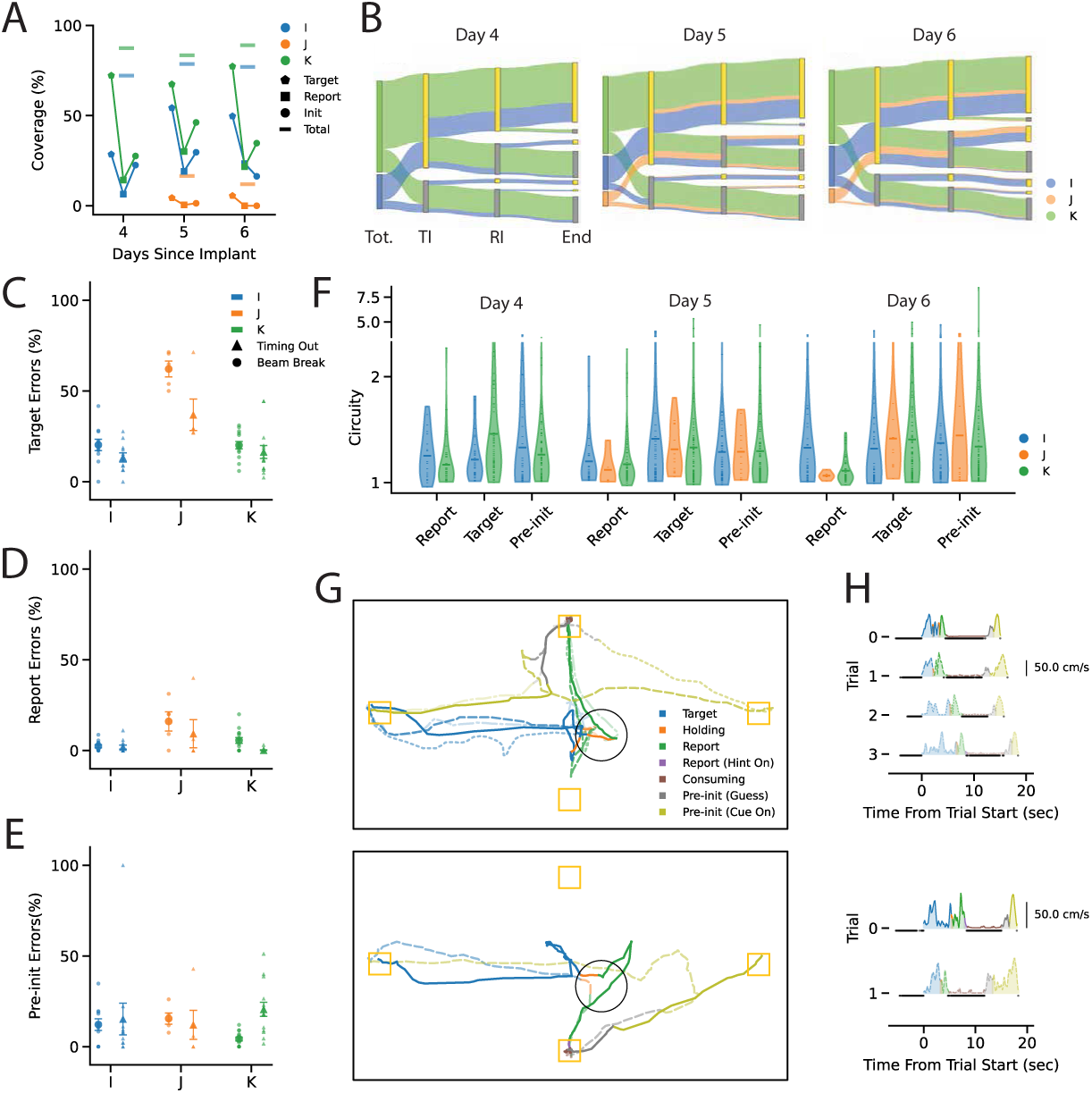
Behavioral performance and errors across task phases. A) Coverage values as computed in (Fig. 2E, K, O) for all animals and days. Colors correspond to individual animals. B) Trial phase outcomes by animal for each day. Split ribbons to gold and grey bars represent clean and “with errors” task phase transitions, respectively. To highlight flow into the report phase vertical bars from left to right are: total trials, pre-initiation phase attempts, of each animal (“Tot.”), target phase initiation (“TI”), report phase initiation (“RI”), reward delivered (“End”). C) The percentage of trials with errors during the target phase, color-coded for animals as in (A). Two types of errors are shown: triangles indicate the task timed out errors, circles indicate beam break errors. Small markers indicate individual session averages and large markers indicate the animal means. Error bars are ± SEM of the session averages. D) The percentage of trials with errors during the report phase, color-coded for animals as in (A). Other elements coded as in (C). E) The percentage of trials with errors in the pre-initiation phase, color-coded for animals as in (A). Other elements coded as in (C). F) Distributions of circuity, or route directness, as violin plots. Dots indicate trials, lines indicate the mean for each task phase and the width of the violin represents the density estimate of the data at different values. Animal color codes from (B). G) Example trajectories showing the high similarity of behavior on trials with matching initiation well and matching target location. Each example trial is indicated by a different line style (e.g. dotted vs. solid). All trials were initiated from well 0 with the same target location (black circle). Blue segments are from initiation to target collision (target phase initiation). Orange segments are target hold. Green segments are the report phase, in which the animal moves to the remembered reward location. Brown segments indicate reward delivered and consumption. Grey and yellow segments indicate the post reward sub-phase and cue ON at the next target phase initiation well, respectively. Top panel shows trials where the report was at well 3. Bottom panel shows trials where the report was at well 1. H) Running speed (cm/s) for each trial shown in (G). Color codes from (G).

We implemented two types of error consequences for the animals presented here (rats I, J and K). For animals I and J, errors resulted in an error tone and suspended state penalty of the environment (10 seconds following a beam break error and 100 ms following a timing out error). After serving the penalty, I and J were allowed to continue the trial. For the purposes of analyses that make a distinction between correct trials and incorrect trials, the trial phases in continuation of an error were excluded. For animal K, errors resulted in the trial aborting and a 0.5-1.5 second suspension of the environment, precluding the large report reward following a completed trial. These error contingencies are evident when examining trial phase outcomes. For animal K, no trials that entered an error state were rescued as a success in a later phase (Figure 3B). The target phase tended to have errors more often than the report phase, and in both cases beam break errors were more common (Figure 3C,D). Error percentages in the pre-initiation phase were similar to those in the target phase, except it was timing out errors that were more common for animals I and K (Figure 3E). Animals showed varied amounts of each error type and patterns of errors over a session (Supplemental Figure 1).

During all phases of the task, trajectory circuity was clustered near one, indicating the paths taken were largely direct paths from origin to endpoint (Figure 3F). In addition, common trajectories show qualitatively similar velocity profiles even when they come from different blocks (Figure 3G,H).

One limitation of the implementation of the design presented in this paper is reachability of target locations. Due to the need for modularity with another experiment utilizing this setup, walls were placed outside of the areas of the floor held up by support material. The 80/20 can be seen in the overhead view around the border of the environment due to its high IR reflectance (Figure 1A). For this reason, the targets, which are being projected from below, could not be projected to the edge of the explorable environment. Furthermore, rats tended to interact on the inside aspect of the targets, the side in line with the angle of approach (Figure 2F). This tendency, compounded with the target size and target location, effectively shrank the explored environment by a distance of 1 target radius from the area covered by the projected targets. Future implementations can quite easily avoid these limitations. Moving the boundaries of the environment inside of the area occluded by the support material would avoid having explorable areas of the environment unreachable by the projector. Additionally, target shapes and orientations could be modified to encourage collisions close to environment boundaries.

### Medial Entorhinal Implant Outcomes and Encoding of Task Goals

To illustrate the viability of the data collected during this task, we implanted Neuropixels 1.0 probes into the MEC of rats I, J, K (Supplemental Figure 2). Note that all behavioral data described above are from post-implant sessions. To validate the feasibility of MEC electrophysiological recordings during the behavioral task, we summarized the number of good units per day and computed tuning curves. While longer term implant viability was variable, the first few days post implant recovery had ample neurons (Figure 4A). Example cells with head direction tuning and others with high spatial stability are among the population of cells recorded, as would be expected in MEC (Figure 4B,C). We report head direction tuning sharpness and stability through the distribution of the mean vector lengths (MVLs) and the Pearson correlation coefficients (r) for each cell (Figure 4D). Additionally, we report the distribution of spatial stabilities with the distribution of Pearson correlation coefficients (Figure 4E). The population of neurons as a whole from each session were a qualitative match with other published MEC results (Hafting et al. 2005; Sargolini et al. 2006; Solstad et al. 2008; Kropff et al. 2015).

**Figure 4.**
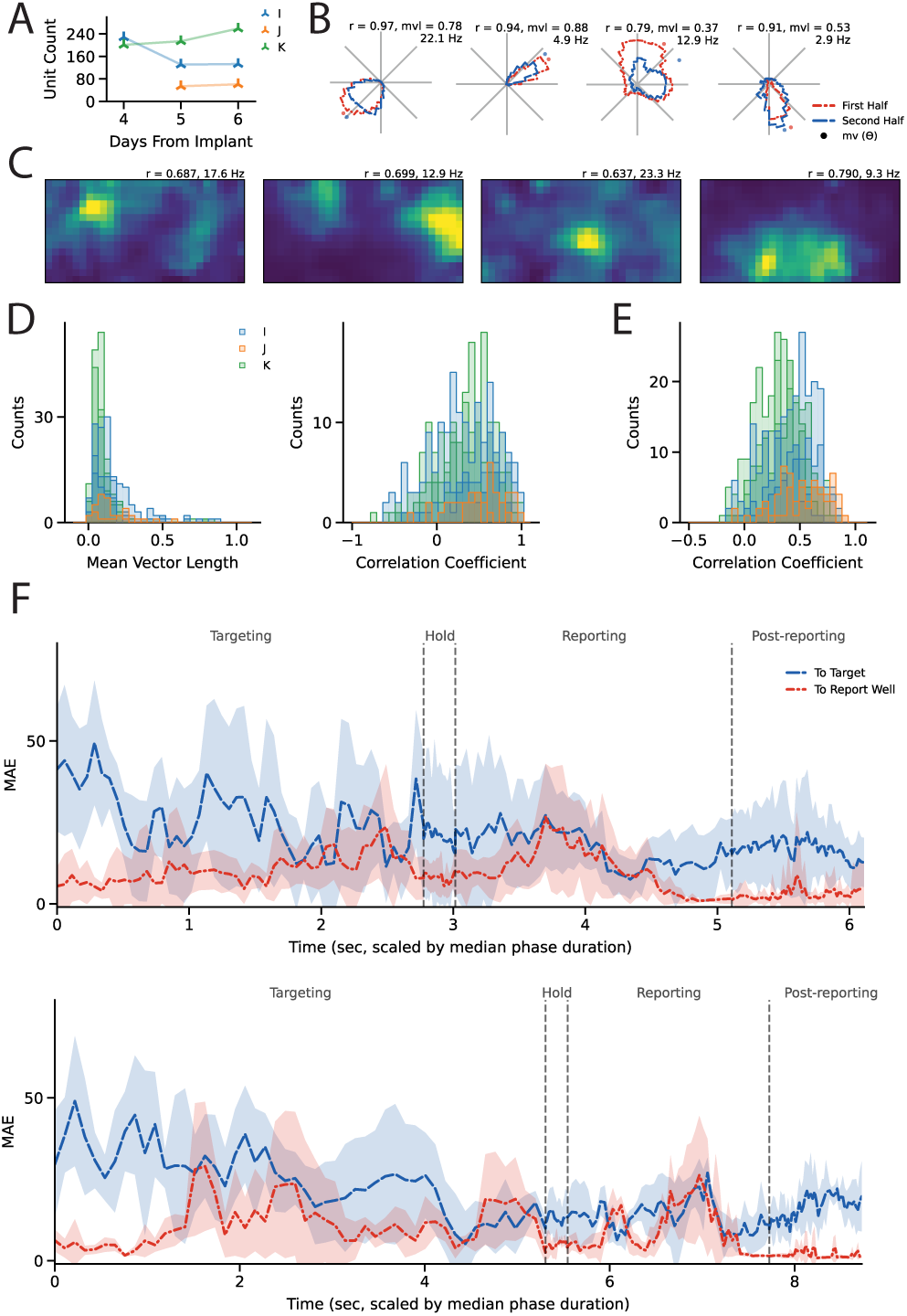
Illustrating neural data viability. A) Unit count over days for each animal. Color coded by animal. B) Four example putative head direction cell tuning curves. Blue and red tuning curves are computed from the first half and second half of a session. The Pearson correlation coefficient (r) between the first-half and second-half head direction tuning curves, mean vector length (mvl), and peak firing rate (Hz) are reported above the tuning curve for each unit. C) Four example putative spatial cell tuning curves. The Pearson correlation coefficient (r) between tuning curves computed from the first-half and second-half of a session, and maximum firing rate bin (Hz) reported for each unit. Color coded for minimum (blue) and maximum (yellow) values. D) Histograms of cells recorded from each animal and recording day. Histograms of mean vector lengths (left) and histograms of the Pearson correlation coefficients (right) as in (B). Color coded by animal as in (A). E) Histograms of correlation coefficients of spatial maps as in (C). F) Multi-layer perceptron (MLP) models trained to predict the distance to target (blue) or the distance to report well (red). Data are shown for the two highest coverage days. Lines show the average model accuracy, mean absolute error (MAE), within a trial. Shading is ± the standard deviation. To accommodate phases of different absolute lengths each phase is first mapped to percent progress and then interpolated. After being combined across trials, the MAE trajectory for each phase is mapped onto its median duration.

To examine some of the task-specific coding that may be captured by the population activity, we trained multi-layer perceptrons (MLPs) to predict the distance from the animal to the “target location” or “report well location” with the preceding 400 ms of neural data. We found that the model’s decoding error for distance to the report well was low throughout the trial, with a notable drop in accuracy in the middle of the report phase. Accuracy of the predicted distance to target shows an improving trend as the target phase progresses (Figure 4F). The notably low decoding error at the end of the trial may reflect the model capitalizing on variables consistently present at that phase, such as behaviors associated with reward consumption, rather than true distance encoding. Together, these results align with the block design of the task: the report location stays constant throughout a block whereas the target is randomly placed on the arena at the beginning of the target phase. Moreover, these findings raise the possibility that MEC population activity can encode both stable and changing task goals across different trial phases, potentially with distinct temporal dynamics for intermediate transient goals versus static reward locations.

## Discussion

This work establishes the viability of an experimental setup that combines some of the powerful features of freely moving behavior with some of the control available in head-fixed virtual reality paradigms. We were able to train rats to perform tasks in the semi-virtual environment, consistent with the ability of prior projection based systems to train rats to perform visual discrimination tasks (Furtak et al. 2009; Jacobson et al. 2014). Our implementation extends this approach by supporting the combination of unconstrained trajectories, trial-structure and experimenter control over task variables. We showed that this technique can be used to promote coverage of an environment through the use of experimenter controlled goal directed trajectories without experimenter interaction. Further, we noted that minor adjustments to the setup would improve instructed coverage.

This behavioral and hardware platform is well positioned to advance the understanding of how internal states or task-dependent goals influence spatial representations in MEC. As the behavioral platform can facilitate the inclusion of different task phases and memory demands, it provides a framework for studying navigational neural activity as it relates to sensory cues, internal goals and task demands. For example, future work could leverage this platform to examine how MEC representations differ between cued and memory guided trajectories or how population activity changes or remains stable during different task phases.

There are many extensions available under the core setup described here. Minor experimental apparatus extensions such as an under floor camera to track locomotor gait and depth cameras to track pose would provide greater access to the richness of behavior. In addition, the task presented here represents a simple design language to probe navigational behavior. Hidden target zones, mobile target zones, or sequential targets are all possible as extensions and would enable experiments aimed at examining prediction or planning. Additional task layers could also be included, such as probabilistic reward delivery, enabling investigation on how uncertainty impacts navigational neural codes.

This design also enabled an initial finding that MEC contains information about the distance to task relevant objects and locations. Additionally, the network may hold consistent representations for broadly relevant locations (e.g. the reward well) while momentarily relevant locations can be held as a reference dynamically.

Taken together, this platform supports the combination of large-scale electrophysiological recordings with controlled yet flexible behavior. Approaches like this, and others (Lester et al. 2025; Lai et al. 2024; Furtak et al. 2009; Jacobson et al. 2014), stand to bridge the gap between naturalistic navigation and experimental control, enabling future work to examine how neural activity supports flexible, goal-directed navigation in complex environments.

## Acknowledgments

We are grateful to Adriana Diaz for help with, and advice on, animal care and histology. We thank Mehdi Azabou for guidance on the MLP analysis and Sabar Dasgupta, Sarah Abdalla, and Paul Nuyujukian for guidance on LiCoRICE. We thank Eshed Margalit for discussions on presenting the MLP analysis. LMG is an HHMI Investigator. This work was supported by the National Institutes of Health 1R01MH126904-01A1 (LMG), R01MH130452 (LMG), BRAIN Initiative U19NS118284 (LMG), The Vallee Foundation (LMG), The James S. McDonnell Foundation (LMG), The Simons Foundation 542987SPI (LMG), and The Howard Hughes Medical Institute (LMG).

## Author contributions

Conceptualization, T.G.F., M.S., L.M.G.; methodology, T.G.F., M.S., A.G.; investigation, T.G.F., X.C.; visualization, T.G.F.; funding acquisition, L.M.G.; project administration and supervision, M.S., L.M.G.; writing – original draft, T.G.F., L.M.G.; writing – review and editing, T.G.F, M.S., A.G., X.C, L.M.G.

## Methods

### Subjects and housing

All procedures were approved by the Institutional Animal Care and Use Committee at Stanford University School of Medicine. Three male Long-Evans rats (aged 6 to 7 months, weighing 414 to 470 grams at implantation) were obtained from Charles River Laboratory. Rats were always given free access to water. Rats were mildly food deprived to 85% of their free feeding body weight and housed on a 12-hour light-dark cycle. Experiments were performed in the light phase.

### In vivo stereotactic survival surgery

Surgeries were performed in a designated laboratory space using aseptic technique. Anesthesia was induced by inhalation of isoflurane gas (2-3%) in an enclosed chamber. During surgery isoflurane gas was maintained between 1-3%, delivered in oxygen, and heat was provided using a self-regulating heating pad. Pre-operative analgesia was administered with buprenorphine (0.4 mg/kg) and ophthalmic ointment was applied to prevent corneal drying. Fur was removed from the scalp, the head was secured using earbars and stereotaxically aligned (Kopf Instruments). Swabs of iodine and ethanol were used to sterilize the scalp and a midline incision was performed to expose the skull after injecting lidocaine along the midline. The skull surface was cleared of soft tissue and dried. A craniotomy was then performed and a ground screw was inserted. Animals were implanted in the left hemisphere 0.5 mm anterior to the transverse sinus (∼8.5 mm posterior to bregma), and 4.1-5.5 mm lateral from the midline with a Neuropixels 1.0 probe at 17-19° posterior from vertical, surrounded by an implant designed for chronic recordings. The ground screw was placed contralaterally above the cerebellum. The skull was coated with Metabond and covered with dental cement to secure all of the components.

Following the surgery, rats received subcutaneous saline for hydration and were allowed to recover on a heated pad. Animals were closely monitored until they were ambulatory and then checked at regular intervals. Postoperative care included the administration of analgesics and when needed, antibiotics and anti-inflammatory agents for several days. After implantation, rats were housed separately and allowed free access to soft food for 1-2 days after surgery. Rats were allowed to recover for 2 days or until the animal was fully recovered prior to the start of electrophysiological recordings.

After recordings were concluded animals were anesthetized, the implant was retracted, and they were euthanized with an overdose of pentobarbital.

### Histology and electrode localization

After explant and euthanasia animals were transcardially perfused with saline and 4% paraformaldehyde. After 24 hours in 4% paraformaldehyde brains were placed in 30% sucrose solution for 72 hours. Brains were frozen and cut into 50-100 μm sagittal sections mounted on slides and stained with DAPI. Recording sites were determined using HERBS (Fuglstad et al. 2023).

### Implant Assembly

Implant parts were printed on a Formlabs stereolithography printer in black resin based on a design from Luo et al. (2020). Probes were mounted to the internal element of the implant and then sharpened on a Narishige grinder. Before gluing up the external elements of the implant around the internal element it was test-fitted and shimmed to verify that the probe was in line with the rest of the implant. The external element was coated with Metabond and Purelube was melted into the implant from the bottom to prevent fluid from wicking up after implanting. Finally, probes were coated with DiD fluorescent dye.

### Arena Details

The arena was built from custom cut aluminum extrusion (80/20). The floor of the arena was 57”x30”x1/4” acrylic with a satin finish. The walls of the arena were 1/4” black matte acrylic 18” high. The wall above reward well 3 had a white cue card. The arena was lit for the overhead camera with three infrared light sources. The floor targets were generated by a JMGO O1 1080p ultra-short-throw LED projector mounted under the floor. A Basler ace ac780-75gc GigE camera with infrared filter removed was positioned above the arena mounted with a wide-angle non-distorting lens.

The wells were printed from standard white resin with a Formlabs stereolithography printer. We modified the well design from (Saathoff et al. 2025) and improved their robustness to urine: moving the printed circuit board out and away from the inside of the well and waterproofing them using epoxy and silicone caulking. Thick-walled Tygon polyvinyl chloride (Tygon PVC) tubing, 1/8” ID, 1/4” OD, connected the inlet of each well to a dedicated NE-500 syringe pump from https://www.syringepump.com/. The outlet of each well was connected to a shared catch vessel under a light vacuum pulled by the building.

### Motorized Slip Ring Details

The slip ring was mounted to the ceiling in the experimental room. Design details are available at https://github.com/giocomolab/turn_up.

### Task Computer and Task Control Hardware

A single Asus Prime z390-A with a GeForce RTX 2070 Super graphics card operated as the task computer. It ran the LiCoRICE model, performed inference with DeepLabCut Live, and stored all task signals and videos. More details about LiCoRICE can be found at https://bil.stanford.edu/licorice. Input and Output from the task computer was accomplished via two parallel ports wired through transistor arrays to allow the task computer to switch a higher current power supply. The line printer Linux kernel device driver was disabled allowing each pin of the parallel ports to be controlled individually as either read or write. The task computer produced a synchronization signal that was passed to a National Instruments DAQ allowing the task signals to be temporally aligned to the neural activity at analysis time.

### Behavioral Training

Animals were trained by building up to the final task. First, during handling, animals acquired familiarity with the evaporated milk as a reward. After initial exposure to the experimental rig, the animals learned the reward wells through the first build-up task: seek reward from three active wells with LEDs on, and avoid one inactive well with its LED off. A beam break on the inactive well resulted in an error tone and a temporary suspension of the task. This helped the animals associate an LED on well with a guaranteed reward. The ability to reach near perfect accuracy without over-training was critical. This task established a fundamental piece of “experimental rig vocabulary” while still allowing for later shaping the animal towards LED off memory guided well visits. Next, animals learned an impoverished version of the final task: targets appeared in either of only two locations, and the task progressed directly after collision (no target hold required). In this build-up task, the reward well blinked its LED every time the report phase was triggered. Additionally, in this build-up task, the targets started very large, nearly spanning the short dimension of the arena, progressively shrinking every time the animal successfully collided. Once the animals showed clear signs of deliberately moving to the floor target, they were advanced to a stage with a fixed target size where they were shaped to hold longer and where the number of forced hint, or report well blinking, trials was decreased.

### Recording navigational information

All data presented here were collected during task time. Body keypoints were extracted using DeepLabCut (Nath et al. 2019). Speed was estimated with first-order adaptive windowing (FOAW) (Janabi-Sharifi et al. 2000). Invalid positions, where speed was not between 2 cm/s and 200 cm/s, were removed from analysis.

Head direction was computed between two keypoints on the head of the animal. Head directions were computed for all times where the speeds of both fiducials were valid.

### In vivo single unit recording and offline spike sorting

Neural data were recorded from Neuropixels 1.0 probes using SpikeGLX acquisition software and National Instruments hardware. Clusters were sorted, separated, and merged into putative single units using Kilosort 2.5 (Pachitariu et al. 2024) and Phy (https://github.com/cortex-lab/phy).

### Circuity

Circuity is the ratio of distance traveled over total displacement. To compute distance traveled the speed (from FOAW velocity) of a trajectory was integrated using the composite trapezoid rule. This numerical method of integration is an approximation of distance traveled. The imperfectness is apparent in 14 of the 1482 phase trajectories appearing to have a circuity of just under one. Total displacement was taken to be the distance between the starting position and ending position of a trajectory.

### Calculation of tuning curves

Spatial tuning curves were generated by first computing an animal position (occupancy) histogram with 5cm x 5cm bins. Then a spike position histogram was generated into the same spatial bins. After first smoothing both the spike position histogram and the animal position histogram using a quasi-Gaussian kernel, the smoothed spike position histogram was divided by the smoothed occupancy histogram to produce the final tuning curve (Butler et al. 2019). Head direction tuning curves were generated using the same quasi-Gaussian kernel smoothing procedure, but with data binned into a polar histogram with 10° bins.

### Stability

Stability was computed as the Pearson correlation between tuning curves generated from the first-half and second-half of a session. For any neuron appearing in more than one session, the stability computed from the session with the highest spatial coverage was used.

### Multi-layer Perceptron

We first downsampled by rejecting error trials and trials with phases that were outliers in duration. To determine outliers we formed a linear boundary for each phase (slope of distance to target over target phase duration, and slope of distance to report over report phase duration) chosen manually, to separate long in duration outlier phases (Supplemental Figure 3). Trials were then split into the training set (60%), validation set (20%), or test set (20%). Within trials, spikes were assigned to contiguous bins 20 ms in length, the sampling period of the position data. The goal of the model was to predict the distance to the target location or reward well location using a context window of 400 ms into the past. Each trial was split into overlapping samples 400 ms in length for all anchor points separated by 20 ms.

To prevent model over-fitting on segments of data where we did not expect accurate predictions, we implemented a heuristic weighting scheme. This scheme asks the model to prioritize learning from putatively high signal periods. Each sample was weighted between zero and one (full weight) based on where the sample anchor point falls relative to trial phase transitions. For the “distance to target” model the heuristic weighting scheme was as follows: (a) samples from before the target phase start, when the animal is consuming the trial initiation reward, were ignored (b) the first second of the target phase, largely when the animal is turning around, was assigned half weight (c) for samples from the target phase where the animal will not arrive at the target in less than six seconds, putative low engagement samples, were assigned half weight (d) samples from the report phase were progressively down weighted from half weight in the first second to twenty hundredths for the remainder of the phase (e) samples during reward consumption were assigned ten hundredths weight. For the “distance to report well” model the heuristic weighting scheme was as follows: (a) samples from before the target phase start were ignored (b) samples within the first second of the target phase were assigned half weight (c) samples from the target phase where the animal will not arrive at the target in less than six seconds, were assigned half weight (d) samples from the report phase were all fully weighted (e) samples during reward consumption were assigned half weight.

Models were evaluated and compared based on their mean absolute error (MAE) on full weight validation set samples. The best model was chosen through a hyperparameter sweep over learning rates (1e-5, 3e-5, 1e-4, 3e-4, 1e-3, 3e-3, 1e-2), hidden layer dimensionality (three layers (hidden dimensions 512, 256, 128), four layers (hidden dimensions 512, 512, 256, 128), and four layers (hidden dimensions 1024, 512, 256, 128)), network unit dropout proportion (0.1, 0.3, 0.6), and weight decay penalties (1e-3, 1e-4, 1e-5).

One challenging aspect to summarizing these results arises from the variability of phase durations, which reflects the self-paced nature of the task and the inherent heterogeneity of distances traveled in each trial. To accommodate for these challenges during visualization, a universal timescale was created. The time within each phase was scaled linearly as progress from phase start to end and interpolated onto a 50-point lattice progressing between zero and one. A phase contributed to the phase averaged error curves in proportion to its duration divided by the median phase duration of that phase. Trial statistics required to generate the ‘median trial’ durations for each phase were computed from the training-set trials. Only trials from the test set were included in the MAE curves.

