## Supplemental Figures for "Structured navigation in a goal-directed task reveals flexible spatial coding"

**Supplemental Figure 1: Coverage, errors and head direction occupancy.**

A) Coverage in the center of the arena. Percent coverage for a square area expanding from the center of the arena.

B) Percent errors over time as a phase progresses (first third, second third, last third) for each animal and day illustrating the relative distribution of errors as a session progresses on average. Triangles show the progression of timing out errors within a session averaged across sessions on that day. Circles show the same progression but for beam break errors. Each line represents the average percentages for all sessions on a day for a single animal.

C) Smoothed head direction occupancy maps for trajectories arriving from each initiation location colored for the initiation well.

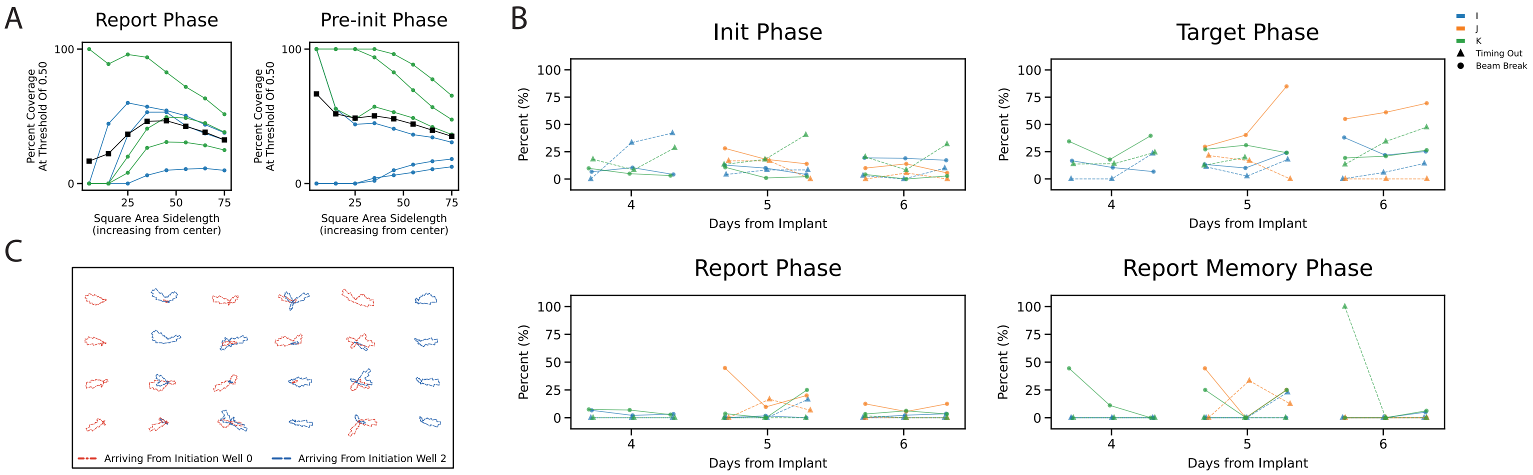
A
Report Phase
Pre-init Phase
Percent Coverage At Threshold Of 0.50
Square Area Sidelength (increasing from center)
B
Init Phase
Target Phase
Report Phase
Report Memory Phase
Percent (%)
Days from Implant
I
J
K
Timing Out
Beam Break
C
Arriving From Initiation Well 0
Arriving From Initiation Well 2

**Supplemental Figure 2: Histological verification of medial entorhinal cortex placement.**

A) Example histology for animal I, left hemisphere. Probe placement can be seen in red.
B) Histology for animal J, as in (A).
C) Histology for animal K, as in (A).

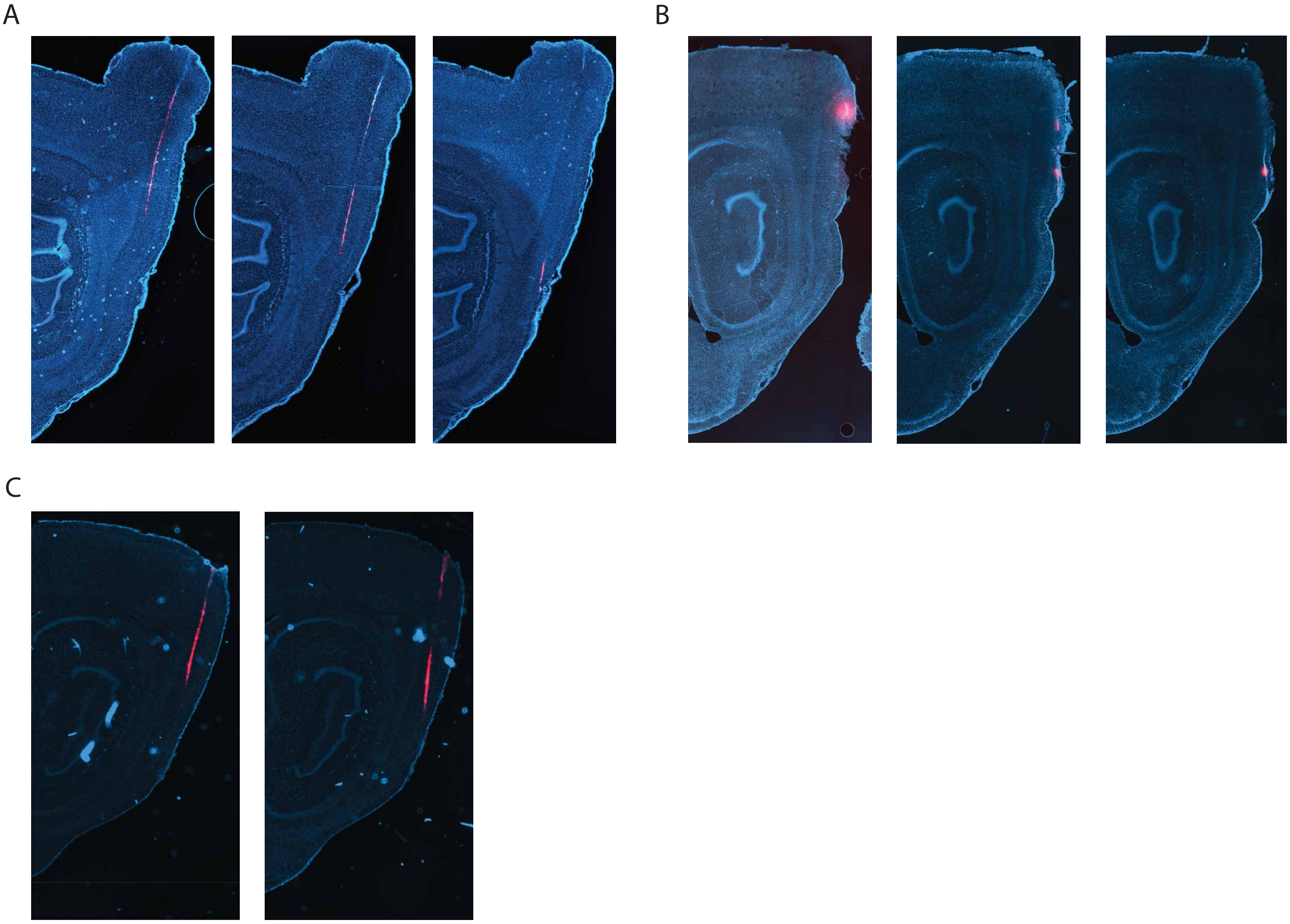
A
B
C

**Supplemental Figure 3: Phase durations**

A) Histograms of target phase durations and report phase durations for animal K on each day.
B) Scatter plot showing each target and report phase from animal K. Green line is the threshold, all phases below this line are dropped from the MLP analysis.

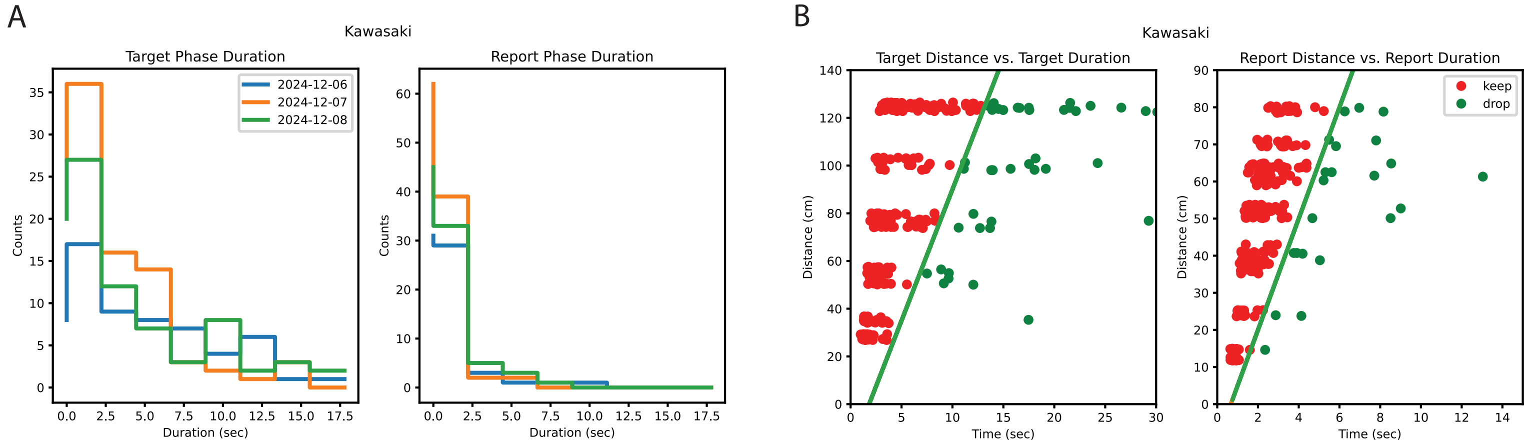
A
Kawasaki
Target Phase Duration
Report Phase Duration
2024-12-06
2024-12-07
2024-12-08
Counts
Duration (sec)
B
Kawasaki
Target Distance vs. Target Duration
Report Distance vs. Report Duration
keep
drop
Distance (cm)
Time (sec)
